# MmpL3, Wag31 and PlrA are involved in coordinating polar growth with peptidoglycan metabolism and nutrient availability

**DOI:** 10.1101/2024.04.29.591792

**Authors:** Neda Habibi Arejan, Desiree R. Czapski, Joseph A. Buonomo, Cara C. Boutte

## Abstract

Cell growth in mycobacteria involves cell wall expansion that is restricted to the cell poles. The DivIVA homolog Wag31 is required for this process, but the molecular mechanism and protein partners of Wag31 have not been described. In this study of *Mycobacterium smegmatis*, we identify a connection between *wag31* and trehalose monomycolate (TMM) transporter *mmpl3* in a suppressor screen, and show that Wag31 and polar regulator PlrA are required for MmpL3’s polar localization. In addition, the localization of PlrA and MmpL3 are responsive to nutrient and energy deprivation and inhibition of peptidoglycan metabolism. We show that inhibition of MmpL3 causes delocalized cell wall metabolism, but does not delocalize MmpL3 itself. We found that cells with an MmpL3 C-terminal truncation, which is defective for localization, have only minor defects in polar growth, but are impaired in their ability to downregulate cell wall metabolism under stress. Our work suggests that, in addition to its established function in TMM transport, MmpL3 has a second function in regulating global cell wall metabolism in response to stress. Our data are consistent with a model in which the presence of TMMs in the periplasm stimulates polar elongation, and in which the connection between Wag31, PlrA and the C-terminus of MmpL3 is involved in detecting and responding to stress in order to coordinate synthesis of the different layers of the mycobacterial cell wall in changing conditions.

## Introduction

Bacterial cell walls are important for shape maintenance and protection from environmental stressors, including antibiotics. The mycobacterial cell wall is distinctive in two ways: first, it has an unusual 3-tiered structure (1), and secondly, it elongates through polar rather than lateral wall expansion (2, 3). These features require careful coordination by numerous enzymes and regulatory proteins. While many of the enzymes that build the cell wall have been described, little is known about how their activity is coordinated to permit cell growth and division, and to maintain the cell wall structure in changing conditions.

The mycobacterial cell wall is made of peptidoglycan (PG), arabinogalactan (AG), and mycolic acids (MA) covalently linked together to form the MA-AG-PG complex (1, 4). The peptidoglycan layer is made from undecaprenyl-disaccharide-pentapeptide (lipid II) precursors which are flipped from the cytoplasm to the periplasm by flippase MurJ/ MviN before incorporation into the peptidoglycan wall by transglycosylases and transpeptidases. Arabinogalactan is composed of a galactan stem with arabinan branches (5) and is important for cell wall integrity and for anchoring the mycolic acid layer to the peptidoglycan (6). The mycolic acids form a lipid bilayer. In the inner leaflet, many of the mycolic acid lipids are covalently attached to the arabinogalactan layer by ester linkages (7). The outer leaflet is made of free lipids, including free mycolic acids, trehalose dimycolates (TDM), trehalose monomycolates (TMM), other glycolipids, and phospholipids (8). Permeability of the cell wall is greatly restricted by the mycolic acid layer because of its tight packing of lipids, thickness, and hydrophobicity (7).

In mycobacteria, the poles are the sites of new cell wall incorporation (2). The poles in mycobacteria are functionally distinct: after septation, the older pole elongates continuously, but there is a pause before elongation begins at the new pole (9). This produces daughter cells of unequal length and physiology after division (9–12). Mechanisms of polar growth are not well understood, though in Actinobacteria DivIVA homologs are critical for this process. In mycobacteria, Wag31 (AKA DivIVA, MSMEG_4217, Rv2145c) is essential for localizing synthesis to the poles, but it is not clear how. Based on the elongasome model in lateral growers, the assumption was that Wag31 recruits peptidoglycan transglycosylase enzymes to the site of elongation. This model works in *Corynebacterium*, where the DivIVA homolog recruits RodA to the pole (13). However, this model is not applicable in mycobacteria, as the transglycosylases do not localize to the poles (14). Point mutants of Wag31 had altered vancomycin staining at the cell poles, which suggested that Wag31 regulates peptidoglycan (15). However, this study did not control for Wag31 protein levels, or differences in permeability to vancomycin, making it difficult to make conclusions about a direct role of Wag31 in peptidoglycan metabolism.

There is no evidence that Wag31 directly regulates any factors in the peptidoglycan synthesis pathway. Co-immunoprecipitation studies indicate Wag31 interacts with fatty acid synthase enzymes (16, 17), MmpL3 and other factors involved lipid metabolism (16). While it does not co- localize with any enzymes in peptidoglycan metabolism, Wag31 does co-localize with MmpL3, the flippase for trehalose mono-mycolates, and other enzymes in mycolic acid synthesis pathway (18, 19). Wag31 also co-localizes with PlrA, a putative regulatory protein which has the same depletion phenotype as Wag31, resulting in short, bulged cells with delocalized peptidoglycan metabolism (20).

MmpL3 (MSMEG_0250, Rv0206c) is an essential transporter of trehalose monomycolates (TMM) from the cytoplasm to the periplasm. MmpL3 belongs to the RND transporter superfamily which powers transport of substrates via proton motive force (21–23). MmpL3 has a large cytoplasmic C-terminal domain which is required for its localization at the poles and interaction with some other proteins, but is dispensable for growth (24–26) MmpL3 is phosphorylated by PknB and dephosphorylated by PstP though the function of this phosphorylation has not been demonstrated (27).

In this work we show that Wag31, PlrA and MmpL3 have connected functions in polar growth, and that MmpL3’s recruitment to the poles is dependent on Wag31 and PlrA. We also show that Wag31, PlrA and MmpL3 all de-localize from the pole in starvation stress; thus, their localization correlates with changes in cell wall metabolism. We found that the localization of MmpL3 is sensitive to perturbations of peptidoglycan metabolism but not inhibition of MmpL3, while inhibition of MmpL3 de-localizes both peptidoglycan and mycolic acid metabolism. Our data show that de-localization of MmpL3 through truncation of its C-terminal domain results in morphological changes, defective responses to starvation, and decreased elongation at the old pole, but doesn’t result in de-localized cell wall metabolism. Our data support a model in which transport of TMMs activates polar peptidoglycan and mycolic acid metabolism irrespective of their location, and Wag31, PlrA and the C-terminus of MmpL3 help the cell sense and respond to environmental changes and coordinate synthesis of the 3-tiered cell wall.

## Results

In an effort to better understand the molecular function of Wag31, we looked for factors that were genetically linked to *wag31* by conducting a suppressor screen. We had previously generated a number of *wag31* partial loss of function point mutants (28). We passaged several slow-growing *wag31* point mutants - D7A, K20A, and L34A - for several days, struck them out regularly and looked for large colony mutants. We then isolated fast-growing suppressors and used whole genome sequencing to map the suppressor mutations (Fig. 1A, S1B). We identified numerous point mutations in separate suppressor strains in the *MSMEG_0251* gene, which is just upstream of *mmpl3* in an operon, and which contains the promoter for *mmpl3* (Fig. 1A) (29). Some of the suppressor mutants would cause loss of function of MSMEG_0251, but some were only point mutations. Many of the mutations we found were near the transcriptional start sites of *mmpl3*.

**Figure 1.**
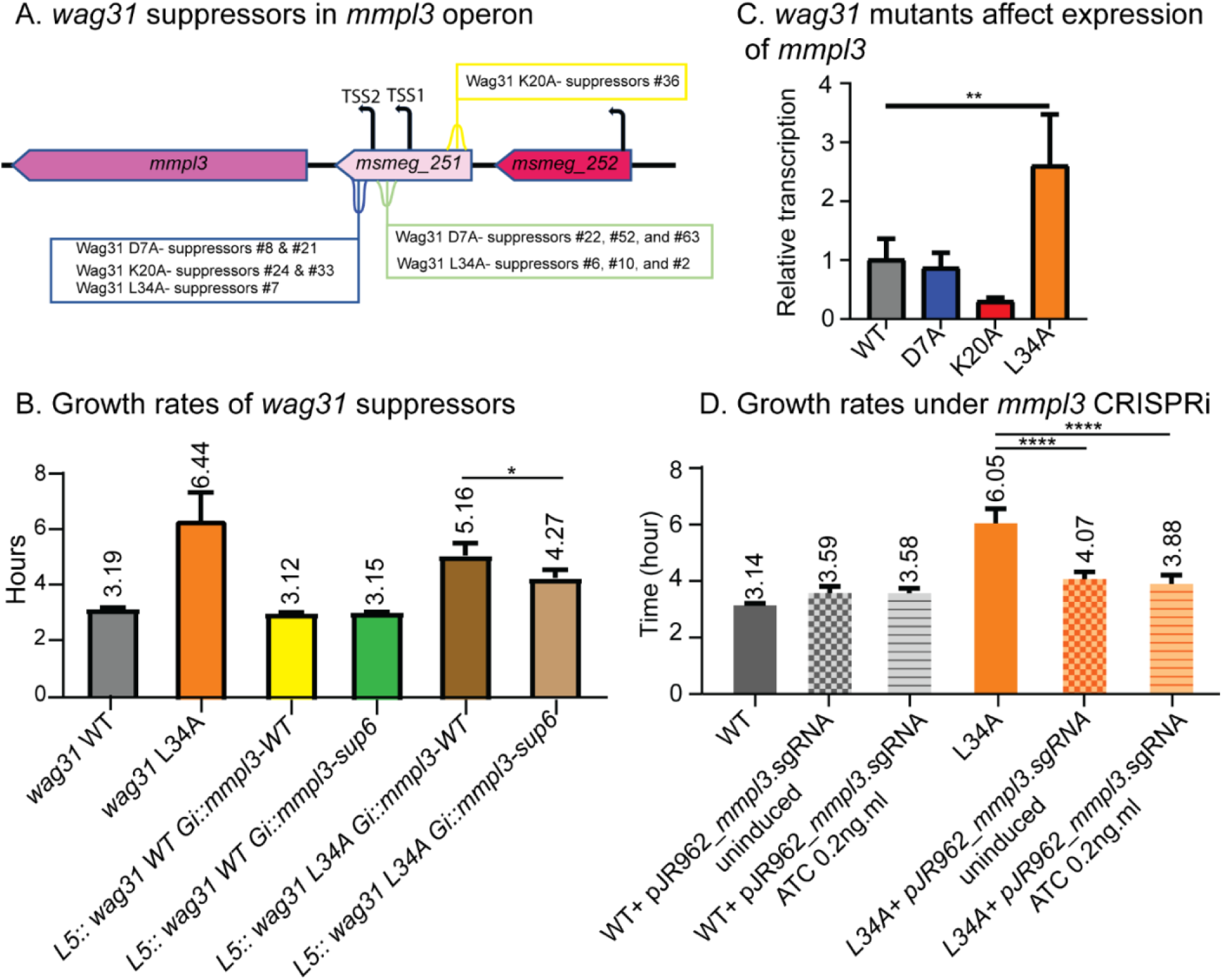
*wag31* and *mmpL3* are genetically connected. (A) Diagram of *mmpl3* operon with positions of *wag31* suppressor mutations indicated. TSS = transcriptional start sites, from (Martini *et. al.*, 2019). (B) Doubling times of *Msmeg* cells expressing the suppressor #6 allele of the *MSMEG_0251*-*mmpl3* operon in *wag31* WT and *wag31* L34A backgrounds. Each bar graph represents an average of three biological replicates. Error bars represent standard deviation. (C) Q-RT-PCR of *mmpl3* transcript levels in *wag31* WT, *wag31* D7A, *wag31* K20A, and *wag31* L34A strains. The graph represents relative expression of *mmpl3* normalized to the housekeeping gene *sigA*. (D) Doubling time of *Msmeg* cells suppressing *mmpl3* transcription by CRISPRi in *wag31* WT and *wag31* L34A backgrounds. Three biological replicates of each strain were used for calculating the doubling time. Error bars represent standard deviation. *, P = < 0.05, **, P=< 0.005, ****, P = <0.0001. All *P*-values are calculated by one-way ANOVA, Dunnett’s multiple comparisons test.

To validate that the mutations in *MSMEG_0251* suppress the mutations in *wag31*, we built an *Msmeg* strain where both *wag31 and msmeg-0251-mmpl3* were complemented at the L5 and Giles integration sites, respectively, and deleted from their native loci. Then, we swapped the *MSMEG_0251-mmpl3* wild type allele with a putative suppressor mutant from suppressor #6, and swapped *wag31* wild-type the with mutant allele L34A (30). We performed growth curves to determine if the slow growth of the *wag31* L34A mutant was suppressed by the missense mutation (D127G) in *MSMEG_0251-mmpl3* found in suppressor #6 (Fig. 1B, S1A). We were unable to clone the rest of suppressor mutants due to their toxicity in *E.coli*. In the *wag31* L34A background, the strain with the suppressor #6 allele of *MSMEG_0251-mmpl3* grows faster than the strain with the *MSMEG_0251-mmpl3* wild type allele, which indicates that the mutation in *MSMEG_0251* partially suppresses the *wag31* L34A phenotype (Fig. 1B).

To determine if these mutations were due to loss of function of *MSMEG_0251*, we characterized a knockout strain from the MSRdb (31). MSMEG_0251 has been shown to interact with MmpL3 (26), so we reasoned that if MSMEG_0251 is an important regulator of MmpL3 activity, then its loss might be able to account for the large differences in growth rate we saw in the suppressor strains (Fig. S1). However, the Δ*MSMEG_0251* cells grow normally, although they are slightly short (Fig. S2B, S2C). We observed no defect in starvation survival (Fig. S2D), and insignificant differences in fluorescent D-amino acid (FDAA) staining in the Δ*MSMEG_0251* mutant (Fig. S2E). The mild phenotypes of the Δ*MSMEG_0251* strain suggest that MSMEG_0251 is not a key regulator of polar growth, and therefore it seemed unlikely that the suppression of the *wag31* defects were due to MSMEG_0251 loss of function (Fig. S2).

We then hypothesized that the point mutations in *MSMEG_0251* might suppress the defects of the *wag31* mutants due to altered transcription of *mmpl3*, since the *mmpl3* transcription start sites are within *MSMEG_0251* (29). To determine if the *wag31* mutations affected *mmpl3* transcription, we isolated mRNA from *wag31* WT, *wag31* D7A, *wag31* K20A and *wag31* L34A strains and performed Q-RT-PCR (Fig.1C). Our data show that transcription of *mmpl3* is significantly elevated in at least one of the mutants, *wag31* L34A (Fig.1C). To determine if the perturbation of *mmpl3* transcription is responsible for the slow growth of the *wag31* L34A strain, we mildly repressed *mmpl3* transcription using CRISPRi (31, 32) with no or very low induction. We found that slight reduction of *mmpl3* transcription suppressed the growth rate defect in the *wag31* L34A strain (Fig. 1D). This result confirms a genetic connection between *wag31* and *mmpl3*, and indicates that regulation of MmpL3 activity is connected to Wag31 function.

Wag31 and MmpL3 co-localize at the poles and the septa (18, 19), but a direct interaction or regulatory connection between these proteins has not been described. To determine whether Wag31 and MmpL3 work in the same pathway in regulating polar growth, we characterized the phenotype of *mmpl3*-depleted cells (Fig. 2AB). Our results show that depletion of *mmpl3* causes cells to become short and bulgy (Fig. 2B) and to exhibit delocalized peptidoglycan metabolism (Fig. 2C) as measured by staining with the fluorescent D-amino acid HADA (33). This exact phenotype has also been observed in *wag31* (34) and *plrA* (20) depletion (Fig. 2BC). These data suggest that Wag31, PlrA, and MmpL3 work together in restricting cell wall metabolism to the cell poles. We note that FDAA dyes in mycobacteria are incorporated largely by LD-transpeptidases, and that their staining therefore directly reports on peptidoglycan remodeling, rather than new peptidoglycan metabolism (35). However, because FDAA staining does correlate strongly with new cell wall incorporation (14, 35, 36), and our data below, we use it throughout as a proxy of polar cell wall metabolism.

**Figure 2.**
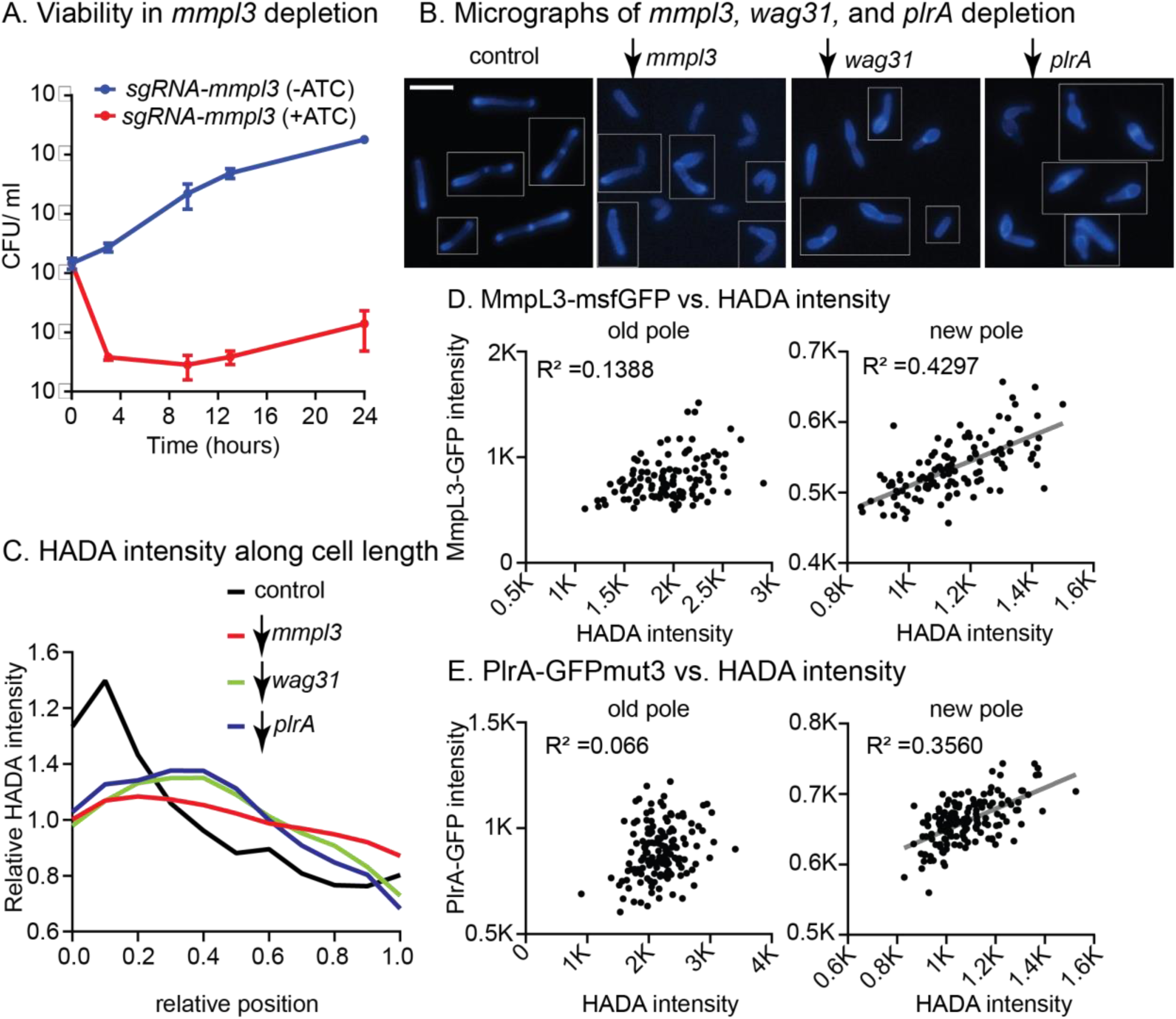
Wag31, PlrA and MmpL3 work in the same pathway to regulate polar growth. (A) Colony Form Units (CFU) of *Msmeg* carrying the *mmpl3* CRISPRi transcriptional repression construct in uninduced (blue, *mmpl3* expressed) and induced (red, *mmpl3* repressed) conditions. (B) Fluorescent micrographs of HADA stained cells in WT (control), and during depletion of *mmpl3, wag31* or *plrA.* Fluorescent signal shows the incorporation of the fluorescent D-amino acid HADA, which reports on peptidoglycan metabolism. The scale bar is 5 microns. (C) Relative averaged HADA intensity across the length of 300+ cells from experiments in B. Data from each cell is oriented so signal from the pole that is brighter by HADA, presumed to the old pole, is at position 0 on the X axis, and the new pole is at position 1. Data are normalized such that the mean of each trace is set to 1, to call attention to differences in distribution of signal. (D) Maximum intensity of MmpL3-msfGFP signal per cell pole in 300+ cells is plotted against the maximum intensity of HADA signal the same pole. Data are separated by pole; the pole of each cell that is brighter by HADA is presumed to be the old pole, and the dimmer pole is presumed to be the new pole. (E) Maximum intensity of PlrA-GFPmut3 signal per cell pole in 300+ cells is plotted against the maximum intensity of HADA signal the same pole, as in D. R2 values in (D) and (E) were calculated by linear regression analysis. The linear fit (gray line) is only shown where there is a correlation.

It has been shown that, after division, Wag31 focus size increases as the new cell pole matures from an elongation-inactive to an elongation-active state (9, 20). We hypothesized that if MmpL3, Wag31 and PlrA work together to regulate cell wall metabolism at the poles, then the size of PlrA and MmpL3 foci might also increase as peptidoglycan metabolism at the new pole increases. We quantified the maximum intensity of MmpL3-msfGFP, PlrA-GFPmut3 and HADA for individual cell poles, and plotted PlrA-GFPmut3 and MmpL3-msfGFP intensities as a function of HADA intensity for each pole. Our data show that there is a moderate correlation between MmpL3-msfGFP and PlrA-GFPmut3 signal with peptidoglycan metabolism at the new poles, but not at old poles (Fig. 2D, 2E). Thus, Wag31, MmpL3, and PlrA accumulate together as the new pole matures, and could be involved in promoting maturation of the new pole into an active growth pole.

Because MmpL3 co-localizes with Wag31 (18, 19), we hypothesized that Wag31 may recruit MmpL3 to the poles. To test this, we depleted *wag31* transcriptionally through CRISPRi (31) and looked at MmpL3-mCherry localization. The *wag31* CRISPRi construct did not allow as rapid depletion as has been seen in other studies (20, 34), so we observed two populations (Fig. 3A): 1) rod-shaped cells that likely have previously-expressed Wag31 at the old poles and 2) short and wide cells which result from more complete Wag31 depletion as in (20, 34, 37). In all cells averaged, MmpL3-mCherry signal is still high at the old poles, likely due to persistence of Wag31 foci there, but is delocalized from new poles, where we would expect no Wag31 to accumulate after induction of the CRISPRi system (Fig. 3B). In cells under 3µm in length, which we conclude have very low Wag31 levels, we observed that MmpL3-mCherry signal was largely delocalized from both poles and re-distributed to the side walls (Fig. 3B). These data indicate MmpL3’s polar localization is dependent on Wag31 during polar elongation.

**Figure 3.**
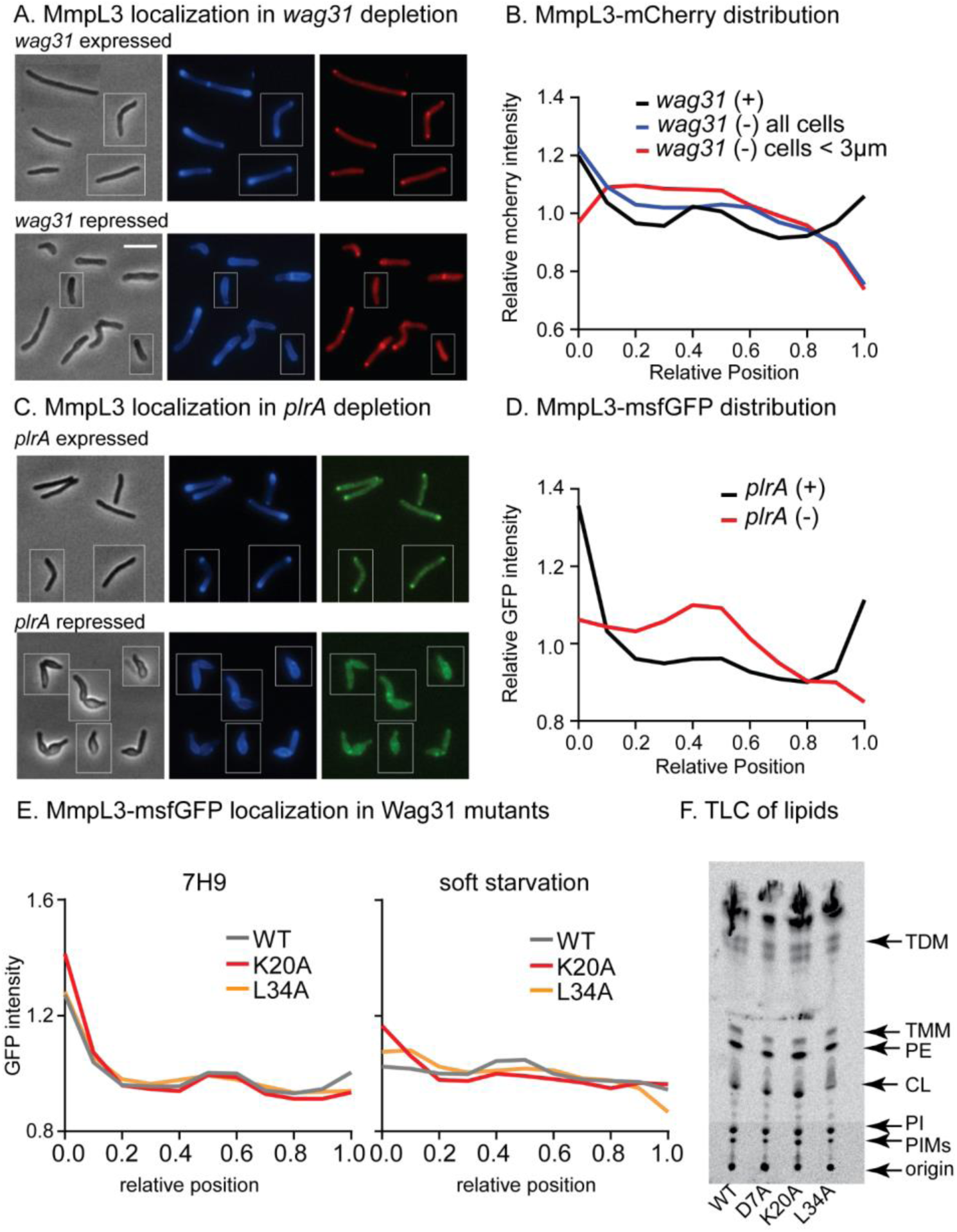
Both Wag31 and PlrA are required for MmpL3’s polar localization. (A) Phase (left) and HADA (middle), Mmpl3-Mcherry (right) images of *Msmeg* expressing MmpL3-mcherry in untreated (top) and CRISPRi repression of *wag31* transcription. Scale bar is 5 microns and applies to all images in figure. (B) Relative averaged MmpL3-mCherry intensity across the length of 300+ cells from experiments in A. Data from each cell is oriented so signal from the pole that is brighter by HADA, presumed to the old pole, is at position 0 on the X axis, and the new pole is at position 1. Data are normalized such that the mean of each trace is set to 1, to call attention to differences in distribution of signal. Relative intensity of alongside of the cell in wag31 depletion. Traces are the average of n=353 *wag31* expressed cells or n=572 *wag31* repressed cells. (C) Phase (left), HADA (middle), MmpL3- msfGFP (right) images of *Msmeg* expressing MmpL3-msfGFP during transcriptional depletion of *plrA*. (D) Relative averaged MmpL3- msfGFP intensity across the length of 300+ cells from experiments in B, as in C. Traces are the average of n=319 *plrA* expressed cells or n=347 *plrA* repressed cells. (E) MmpL3- msfGFP intensity across the length of 300+ cells from strains with the indicated *wag31* alleles. (F) Thin-layer chromatography of C14 labeled lipids extracted from strains with the indicated *wag31* alleles. Lipid species that could be identified are labeled.

As PlrA is also required for polar growth (20), we determined if MmpL3’s polar localization is dependent on PlrA. We transcriptionally depleted PlrA and measured MmpL3-msfGFP distribution (Fig. 3C). Our data show that MmpL3 delocalizes from the poles when PlrA is depleted (Fig. 3D). Thus, polar localization of MmpL3 is dependent on both Wag31 and PlrA.

We hypothesize that Wag31, PlrA and MmpL3 are working together somehow to help coordinate polar cell wall expansion. Previous studies showed that MmpL3 with a truncation of its C-terminal cytoplasmic domain is delocalized from the poles (18, 26) but the cells are still viable, indicating that polar localization of MmpL3 is not essential for growth. To further probe the regulation of MmpL3’s localization, we determined how polar localization of MmpL3 is impacted in *wag31* mutants that are defective for polar growth. We found (Fig. 3E) that MmpL3 was slightly mis-localized in both logarithmic phase and carbon starvation in these strains.

However, our data indicate that polar localization of MmpL3 does not correlate with polar growth, as the *wag31* K20A mutant is defective in old pole elongation (28) but has increased old pole localization of MmpL3, while the L34A mutant is defective in new pole elongation and has decreased new pole localization of MmpL3.

To determine if Wag31’s control of MmpL3’s localization could regulate MmpL3’s transport activity, we extracted C14-labeled lipids from strains with different *wag31* alleles to compare the relative amounts of TMM and TDM: increased relative levels of TMM/TDM are an indication of decreased MmpL3 transport activity (25, 38). The relative amounts of TMM and TDM were not changed in strains with *wag31* alleles with different polar growth and MmpL3 localization patterns (Fig. 3F). These data indicate that mis-localization of MmpL3 does not affect MmpL3 transport activity and suggest that Wag31 does not regulate MmpL3 transport activity. This conclusion is corroborated by previous work which showed that truncation of the C-terminal domain of MmpL3, which causes complete de-localization of MmpL3 away from the poles, had no obvious effect on cell growth (24–26).

Our results indicate that the connection between Wag31, PlrA and MmpL3 at the cell poles does not regulate MmpL3’s transport activity. We hypothesized that this connection could be involved in sensing changes in metabolism and coordinating regulation of other layers of the cell wall at the pole.

Our results suggest that Wag31, PlrA, and MmpL3 are working together to promote polar growth (Fig. 3). Polar growth in mycobacteria only occurs in nutrient-rich and low-stress conditions – in starvation and other stresses, cell wall metabolism is downregulated and re-distributed along the lateral walls (14, 35, 39) (Fig. 4A). If Wag31, PlrA and MmpL3 are critical for polar growth, we would expect their activity and/or distribution should match the distribution of cell wall metabolism in different conditions. Previous work has shown that when *Msmeg* is switched from logarithmic phase to complete carbon starvation, it stops growing and maintains the same cell morphology as in logarithmic phase – we call this “hard starvation”. But, when it is switched to minimal media with Tween80 as a carbon source, polar elongation is downregulated and septation continues for a division cycle or two, leading to very short cells (40, 41) – we call this “soft starvation”. We find that peptidoglycan and mycolic acid metabolism, as measured by HADA and DMN-tre (42) staining, delocalize from the poles and re-distribute to the sidewall and septa in starvation, with slightly more septal cell wall metabolism apparent in soft starvation, especially at earlier time points when reductive division is occurring (Fig. 4A).

**Figure 4.**
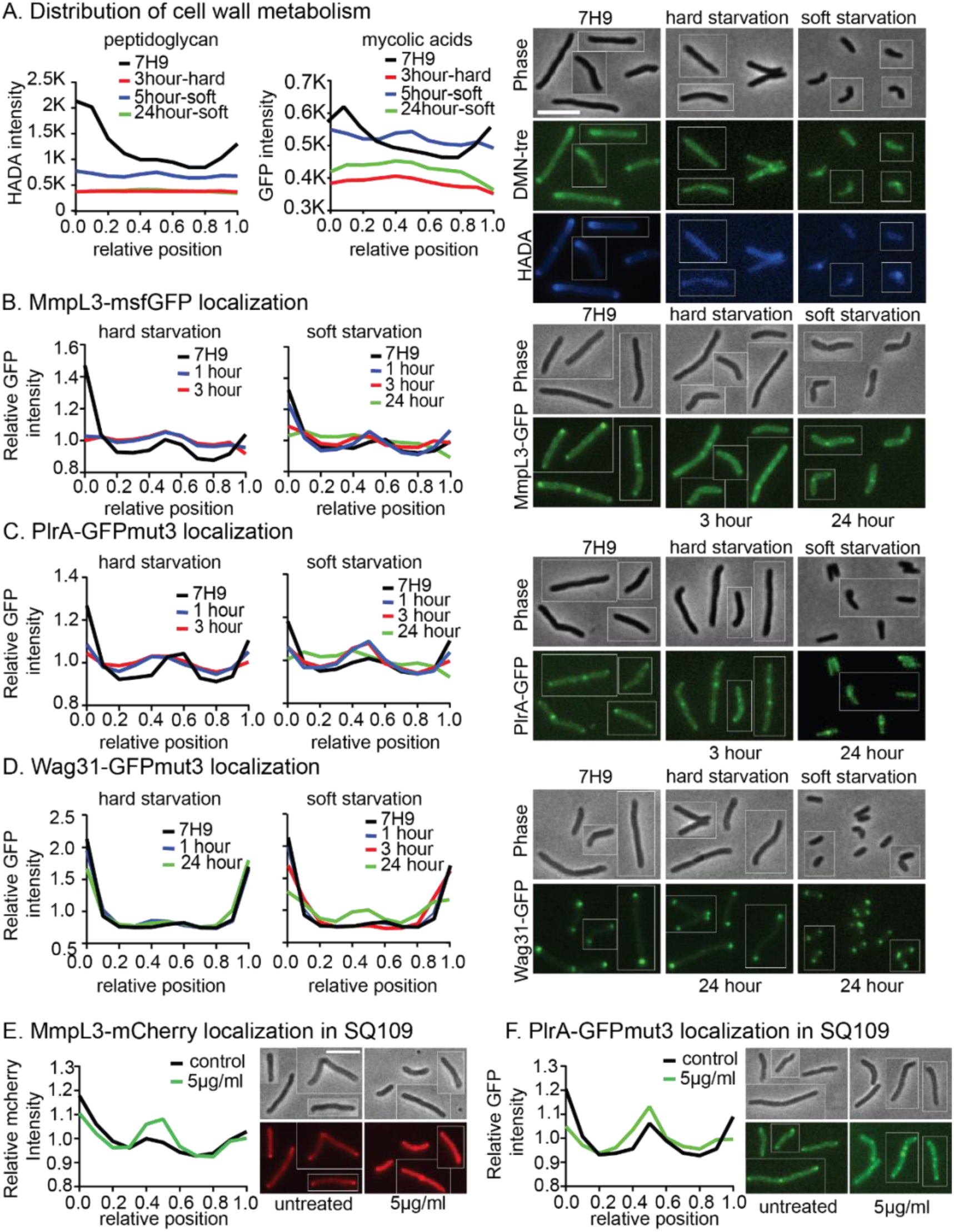
Localization of MmpL3, PlrA and Wag31 responds to energy stresses. (A) Averaged HADA intensity (reporting on peptidoglycan) and DMN-tre intensity (reporting on mycolic acids), in both hard and soft starvation, across the length of 300+ cells, representative micrographs are on the right. Data from each cell is oriented so signal from the pole that is brighter by HADA, presumed to the old pole, is at position 0 on the X axis, and the new pole is at position 1. Analysis was performed on the indicated number of cells, split evenly between three biological replicate cultures: 3 hour hard starvation, n=334; 5 hour soft starvation, n=345; 24 hour soft starvation, n=335; 7H9 control, n=458. (B) Relative, averaged MmpL3-msfGFP intensity in both hard and soft starvation, across the length of 300+ cells, representative micrographs are on the right. Data are oriented as in A. The data analysis was done on three biological replicates of MmpL3-msfGFP strain. Analysis was performed on the indicated number of cells, split evenly between three biological replicate cultures: 1 hour hard starvation, n=328; 3 hour hard starvation, n=314; 1 hour soft starvation, n=369; 3 hour soft starvation, n=366; 24 hour soft starvation, n=420; 7H9 control, n=358. (C) Relative, averaged PlrA-GFPmut3 intensity in both hard and soft starvation, across the length of 300+ cells, representative micrographs are on the right, as in B. Analysis was performed on the indicated number of cells, split evenly between three biological replicate cultures: 1 hour hard starvation, n=382; 3 hour hard starvation, n=355; 1 hour soft starvation, n=351; 3 hour soft starvation, n=423; 24 hour soft starvation, n=906; 7H9 control, n=364. (D) Relative, averaged Wag31-GFPmut3 intensity in both hard and soft starvation, across the length of 300+ cells, representative micrographs are on the right, as in B. Analysis was performed on the indicated number of cells, split evenly between three biological replicate cultures: 1 hour hard starvation, n=333; 24 hour hard starvation, n=428; 1 hour soft starvation, n=352; 3 hour soft starvation, n=349; 24 hour soft starvation, n=967; 7H9 control, n=334. (E) Relative, averaged MmpL3-mCherry intensity across the length of 300+ cells treated or untreated with 5µg/ml of SQ109 for 1hour, data arranged as in B, representative images to the right. (F) Relative, averaged PlrA- GFPmut3 intensity across the length of 300+ cells treated or untreated with 5µg/ml of SQ109 for 1hour, data arranged as in B, representative images to the right.

We investigated whether localization of Wag31, PlrA, and MmpL3 correlate with cell wall metabolism through different conditions (Fig. 4). Our data show that PlrA and Mmpl3 are re- distributed from the poles to the side walls in hard starvation (Fig. 4B, 4C). Although we starved the cells for up to 24 hours, we did not detect changes in Wag31 localization in hard starvation (Fig. 4D). Wag31 forms large, stable oligomers (43, 44) which apparently are not remodeled in abrupt, hard starvation (Fig. 4D). In soft starvation, our analysis reveals that PlrA, and MmpL3 re-localize from the poles to the septa and side walls at early time points and to the sidewalls at later time points. We also find that Wag31 foci become smaller and more septal after prolonged soft starvation, indicating that the Wag31 oligomers are remodeled in this condition (Fig. 4ABCD). These results show that the distribution of PlrA and MmpL3 correlates with cell wall metabolism across nutrient conditions.

To determine if the observed protein re-localization is related to membrane energetics, we treated cells with SQ109. SQ109 is an MmpL3 inhibitor and protonophore, dispersing the proton motive force (PMF) in the inner membrane (38, 45). We found that PlrA and MmpL3 migrate from the poles to the side walls and septa when the cells were treated by SQ109 for an hour (Fig. 4EF). Taken together, these data indicate that the localization of Wag31, PlrA, and MmpL3 is responsive to cell energetics.

SQ109 and starvation have global effects on cell metabolism – we wanted to know if the localization of MmpL3 and PlrA might also be sensitive to more targeted inhibition of the cell wall. We found that MmpL3 and PlrA substantially re-localize from the poles to the side wall in moenomycin, which inhibits the transglycosylase activity of Class A Penicillin Binding Proteins (Fig. 5AB). MmpL3 also delocalizes when the lipidII flippase *murJ* is transcriptionally depleted (Fig. 5C). Together, these data suggest that transport or use of lipidII in the periplasm regulates the localization of MmpL3 and PlrA, and that MmpL3 may be regulated by a component of the peptidoglycan synthesis pathways.

**Figure 5.**
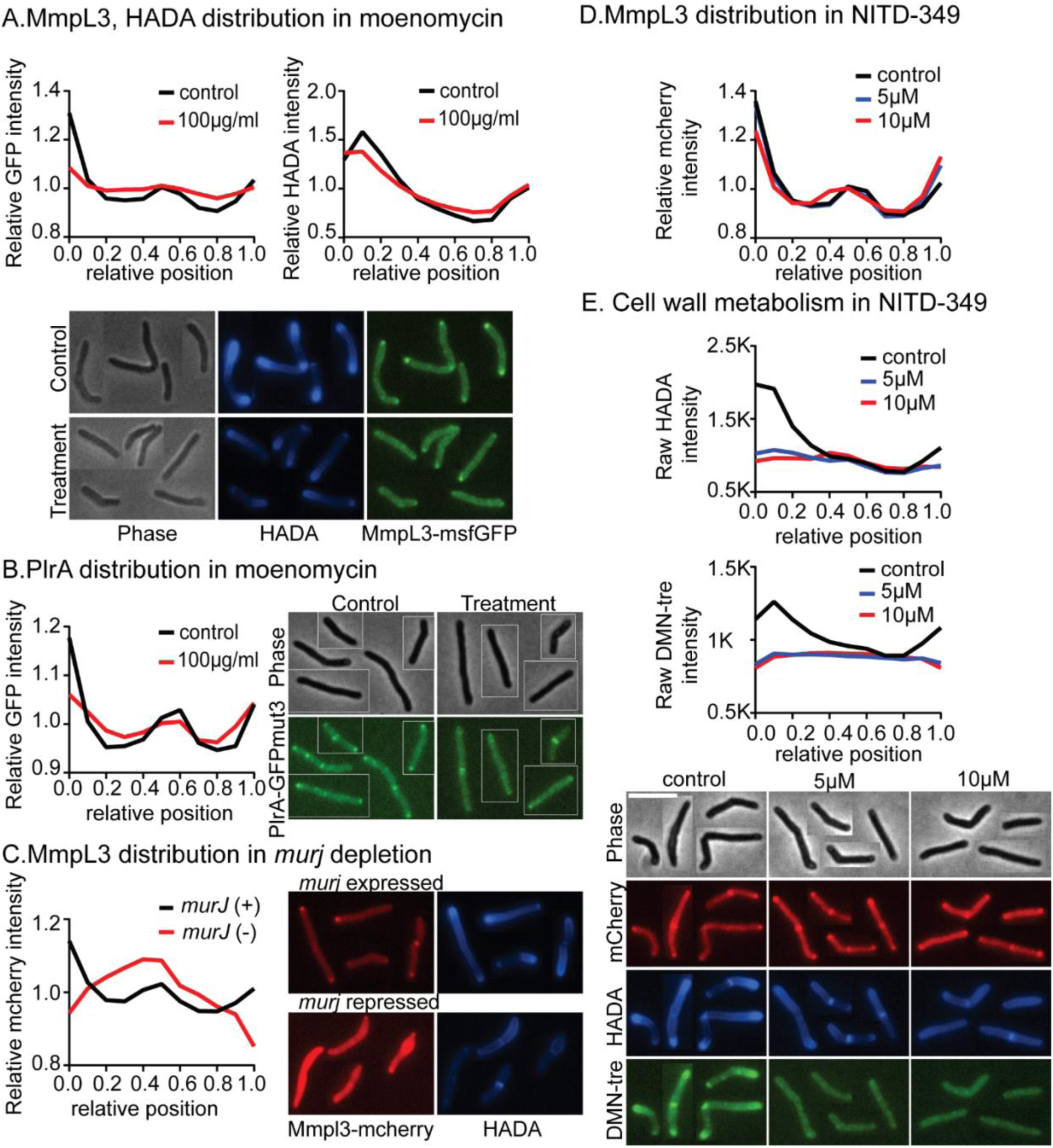
MmpL3 connects regulation of peptidoglycan and mycolic acid metabolism. (A) Relative, averaged HADA and MmpL3-msfGFP intensity across the length of 300+ cells treated or untreated with moenomycin at 100 µg/ml for one hour. Representative micrographs are below. Data from each cell is oriented so signal from the pole that is brighter by HADA, presumed to the old pole, is at position 0 on the X axis, and the new pole is at position 1. (B) Relative, averaged PlrA-GFPmut3 intensity across the length of 300+ cells treated or untreated with moenomycin at 100 µg/ml for one hour. Data are oriented as in A, representative micrographs are to the right. (C) Relative, averaged MmpL3-mCherry intensity across the length of 300+ cells expressing *murJ* (control, black) or repressing *murJ* through CRISPRi for 16 hours (red). Data are oriented as in A, representative micrographs to the right. (D) Relative, averaged MmpL3-mCherry intensity across the length of 300+ cells treated or untreated with NITD-349 at 5 or 10 µg/ml for one hour. Data are oriented as in A, representative micrographs are below E. (E) Averaged HADA and DMN-tre intensity across the length of 300+ cells treated or untreated with NITD-349 at 5 or 10 µg/ml for one hour. Data are oriented as in A, representative micrographs are below.

Next, we investigated whether inhibition of MmpL3 impacts its localization and the distribution of cell wall metabolism. We used the MmpL3 inhibitor NITD-349 for one hour treatments, to disrupt trehalose monomycolate (TMM) transport without affecting the proton motive force (38). MmpL3 remains at the poles when it is inhibited, though it increased at the new pole, and slightly decreased at the old pole (Fig. 5D). But, both peptidoglycan and mycolic acid metabolism is inhibited and delocalized in response to MmpL3 inhibition (Fig. 5E). These data indicate that MmpL3’s localization is not dependent on its activity, but that polar peptidoglycan synthesis is dependent on MmpL3 activity (Fig. 5DE).

MmpL3 has an N-terminal transporter domain, whose function in TMM transport is well established (46), and a poorly-conserved and unstructured C-terminal cytoplasmic domain. The C-terminal domain of MmpL3 is required for its localization at the poles and the septa (47), and previous studies showed no overt phenotypes of the MmpL3 ΔCT strain, suggesting this truncation has little or no effect on MmpL3’s essential function in TMM transport (24, 26). Our data also indicate that MmpL3’s TMM transport role is functionally independent of its localization and connection to Wag31, mediated by the C-terminal domain (Fig. 3EF, 5D). Since MmpL3’s polar localization is dependent on both its C-terminal domain and Wag31 and PlrA, the C-terminal domain must interact, directly or indirectly, with Wag31 or PlrA. We characterized phenotypes of the MmpL3 ΔCT strain (MmpL3_1-749_) (Fig.6) to better understand the function of the C-terminal domain of MmpL3 and the polar recruitment of MmpL3 by Wag31 and PlrA.

**Figure 6.**
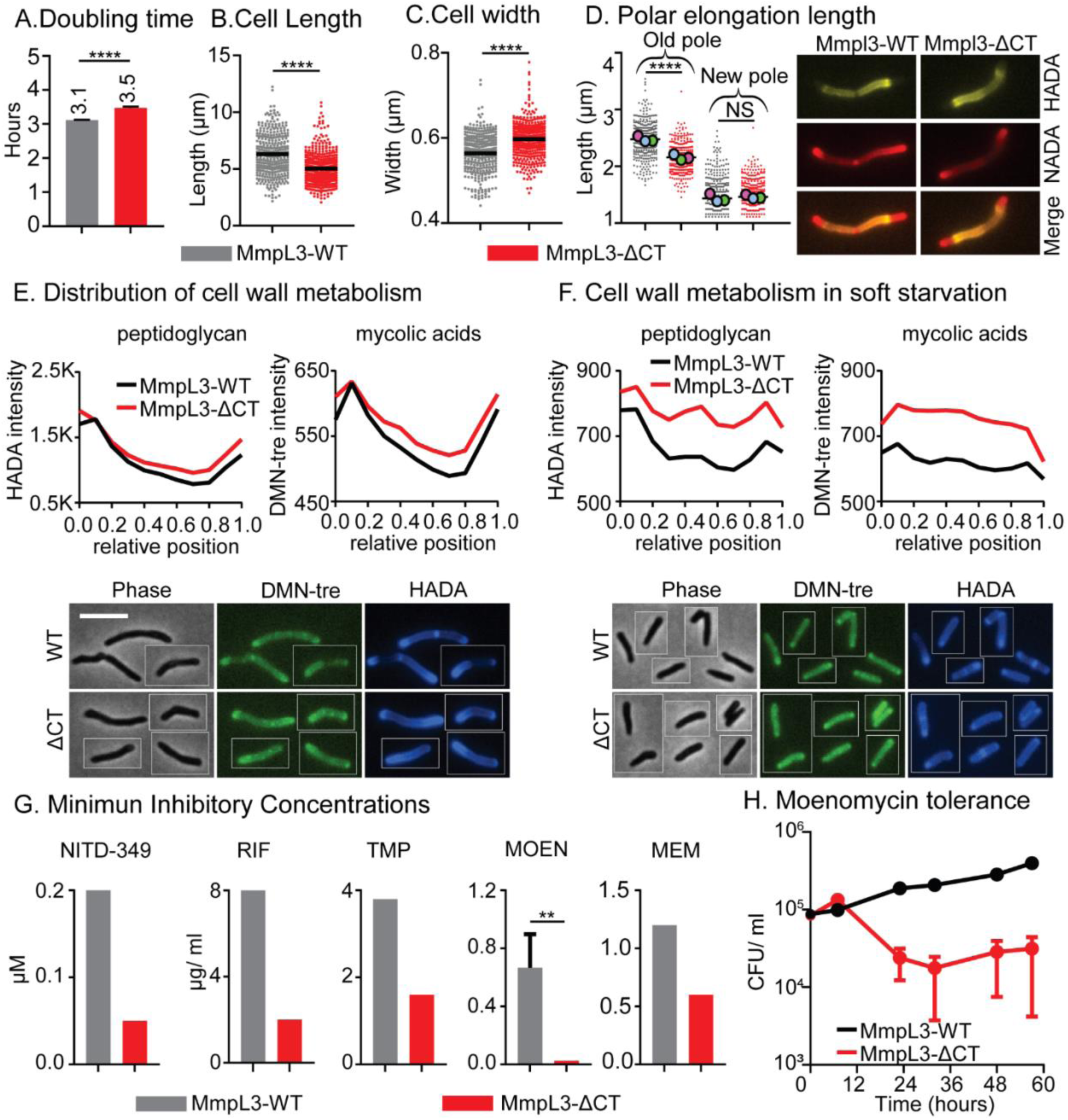
The MmpL3 C-terminal domain regulates polar elongation and starvation response. (A) Doubling time of *Msmeg* with *mmpl3* WT or *mmpl3-ΔCT* alleles. Data represent an average of three biological replicates, error bars represent standard deviation. (B) and (C) The length and width of cells in A. Black bar is the mean. (D) Length of polar elongation in the *mmpl3 WT* and *mmpl3-ΔCT* strains during 1.5 hours of log. phase growth between initial HADA stain and later NADA stain. The length between the pole tip and the end of the HADA signal was measured for each cell pole. Black bar is at the mean, colored balls represent the means of ∼100+ cells from each biological replicate culture. The representative images show the double stained cell, HADA (false colored yellow) and NADA (false colored red). (E) Averaged HADA and DMN-tre intensity across the length of 300+ cells of each strain in log. phase. Data from each cell is oriented so signal from the pole that is brighter by HADA, presumed to the old pole, is at position 0 on the X axis, and the new pole is at position 1. Representative images are below. The scale bar is 5 microns and applies to all images. (F) Averaged HADA and DMN-tre intensity across the length of 300+ cells of each strain after 5 hours of soft starvation, data are oriented as in E, representative images are below. (G) Minimum inhibitory concentrations (MICs) of indicated strains were measured using spot assay on an LB Lennox plates. Three biological replicate cultures of each strain were measured, error bars are not shown when the values were identical between the replicates. Error bars represent standard deviation. NITD-349 is an MmpL3 inhibitor; RIF= rifampin; TMP= trimethoprim; MOEN= moenomycin; MEM= meropenem. (H) CFU of indicated strains during soft starvation and treatment with 100 µg/ml of moenomycin. Error bars represent the standard error of the mean. ns, p >0.05, **, P=< 0.005, ****, P = <0.0001

We found that cells of the MmpL3 ΔCT strain are shorter, wider, and grow slightly slower than the MmpL3 WT cells (Fig. 6A-C). To distinguish whether the short cells result from changes in elongation or cell division, we measured the length of polar elongation in 1.5 hours of growth using sequential FDAA staining (Fig. 6D) and found that MmpL3 ΔCT cells have slower elongation at the old pole (Fig. 6D). Next, we looked at peptidoglycan and mycolic acid metabolism using HADA and DMN-tre (Fig. 6E) and found that staining of both cell wall layers is still polar in the MmpL3 ΔCT strain, though there is a slight increase of staining along the lateral walls (Fig. 6E). These data indicate that the C-terminal domain of MmpL3 is needed for maximal polar elongation at the old pole, but that polar localization of MmpL3 is not required for polar cell wall metabolism.

Due to the localization changes of MmpL3 and PlrA in starvation, we hypothesized the connection between Wag31, PlrA and the C-terminus of MmpL3 is involved in regulating cell wall metabolism in response to environmental changes. To test if MmpL3 ΔCT cells are defective in starvation response, we soft-starved for 5 hours and observed that the MmpL3 ΔCT strains had greater HADA and DMN-tre staining than MmpL3 WT (Fig. 6F). This result indicates that the C-terminal domain of MmpL3 promotes downregulation of cell wall synthesis during stress (Fig. 6F).

To determine if the MmpL3 C-terminal domain affects antibiotic susceptibility during growth, we performed plate-based MIC assays with antibiotics that target both cell wall synthesis enzymes and cytoplasmic enzymes. We found that the MmpL3 ΔCT is more sensitive to all the antibiotics we tested, even those not connected to cell wall metabolism like rifampin and trimethoprim.

From these results, we conclude that the MmpL3 ΔCT strain likely has greater permeability to different classes of antibiotics (Fig. 6G), and therefore the C-terminal domain of MmpL3 promotes antibiotic resistance broadly, likely by controlling cell wall synthesis.

Since downregulation of cell wall metabolism in starvation contributes to antibiotic tolerance (48, 49), we measured how truncation of the C-terminal domain of MmpL3 affected tolerance to moenomycin in carbon starvation. We measured colony forming units (CFU) in moenomycin-treated cultures during soft starvation (Fig. 6H).. Our results show that the MmpL3 ΔCT strain dies faster than MmpL3 WT (Fig. 6G). These data show that the MmpL3 ΔCT strain is more sensitive to moenomycin in starvation as well as growth conditions, which could be due to increased peptidoglycan metabolism in starvation (Fig. 6F) or due to increased permeability.

## Discussion

Our results indicate that one important function of Wag31 is to regulate MmpL3, though we have not established whether this regulation in direct or indirect. We showed that *mmpl3* and *wag31* are genetically connected and *mmpl3* expression is regulated by Wag31 function (Fig. 1). Our results indicate that *wag31*, *plrA* and *mmpl3* are part of the same pathway in regulating polar growth (Fig. 2) and in responding to environmental stresses (Fig. 4,5), and that Wag31 and PlrA recruit MmpL3 to the pole (Fig. 3), and mutants of Wag31 have altered localization of MmpL3 (Fig. 3E). Our characterization of an *mmpl3* mutant that is not localized by Wag31 indicates that Wag31’s regulation of MmpL3 does not affect MmpL3’s transport activity, but promotes downregulation of cell wall metabolism in stress, antibiotic tolerance, and elongation at the old cell pole (Fig. 6). Previous co-localization (19, 47) studies also support the idea that Wag31’s role is more directly connected to mycolic acid metabolism than to peptidoglycan metabolism.

Our data are consistent with a model in which the presence of TMMs in the periplasm promotes polar elongation of all cell wall layers. When we inhibited transport of TMMs into the periplasm with NITD-349, we see delocalized and downregulated peptidoglycan as well as mycolic acid metabolism (Fig. 5E). Delocalization of MmpL3 through truncation of its cytoplasmic domain (47) does not significantly impact the polarity of cell wall metabolism (Fig. 6E), indicating that the location of MmpL3 is not critical for polar growth, but that the transport of TMMs is. We propose that TMMs may interact with other partners at the poles in the periplasm to stimulate polar cell wall synthesis. One possibility is PgfA, which is localized at the poles, interacts with TMMs and is known to be involved in regulating polar cell wall metabolism (50). Previous work showed that MmpL3, LamA and PgfA regulate the asymmetry of polar growth by controlling mycolic acid transport (11, 50). Polar growth requires coordinated assembly of all the layers of the wall; thus, those results indicate that MmpL3, LamA and PgfA are involved in regulating peptidoglycan and arabinogalactan synthesis, directly or indirectly. Our results support the model that MmpL3 activity impacts polar asymmetry and thus must regulate the other layers of the wall. MmpL3 has stronger localization to the fast-growing old pole (Hannebelle et al., 2020; Gupta et al., 2022; Quintanilla et al., 2023). We found that MmpL3 localization correlates with maturation of the new pole (Fig. 2D) and became more symmetric when its activity is inhibited (Fig. 5D). Furthermore, we showed that the MmpL3ΔCT mutant, which cannot be regulated by Wag31, is defective in old pole elongation (Fig. 6D). Our results support the model (Gupta et al., 2022) that polar growth of all cell wall layers is dependent on mycolic acid transport.

We propose that the regulation of MmpL3 by Wag31 and PlrA is important in detecting and responding to stresses, in order to coordinate cell wall synthesis in changing conditions. The polar localization of MmpL3 through Wag31 and PlrA is not significantly regulating MmpL3’s transport activity, since the MmpL3 ΔCT mutant has a minimal growth defect (Fig. 6A), and *wag31* mutants that affect MmpL3 localization (Fig. 3E) do not affect its transport activity (Fig. 3F). While MmpL3 and PlrA are de-localized in response to starvation (Fig. 4BC), disruption of the proton motive force (Fig. 4EF, Fig. S3FGH), and inhibition of peptidoglycan metabolism (5ABC), delocalization of MmpL3 alone is not sufficient to downregulate polar growth, since the MmpL3 ΔCT mutant is completely delocalized from the poles (26, 47) but maintains polarity of cell wall metabolism (Fig. 6E). However, regulation of MmpL3 through its C-terminus, which we presume occurs through Wag31 and PlrA, becomes essential in antibiotic stresses (Fig. 6GH) and helps downregulate cell wall metabolism under starvation (Fig. 6F). Thus, it seems that delocalization of MmpL3 occurs when polar growth is downregulated, but that this delocalization itself does not downregulate MmpL3’s activity, but may instead be an indication that MmpL3 is sensing changes in cell metabolism, and helping downregulate cell wall metabolism globally in response (Fig. 6F), likely through regulatory interactions of its C-terminal domain. One possibility is that MmpL3 could be sensing the status of peptidoglycan metabolism, possibly by detecting lipidII, since lipidIl would be impacted in moenomycin treatment and *murJ* depletion (Fig. 5AC), and since lipidII levels are expected to be delocalized from the polar regions in stress (14, 39).

In this work, we find that Wag31, PlrA, and MmpL3 work together to regulate cell wall metabolism. We show that their cooperation is important for regulating polar growth and polar asymmetry (Fig. 2, Fig. 6D). We propose that periplasmic TMMs are a key signal to stimulate polar peptidoglycan metabolism (Fig. 5E, 6E). Finally, we propose that the C-terminal domain of MmpL3 has a function in sensing and responding to stress through its interactions with Wag31 and PlrA, and that the functioning of the C-terminal regulatory domain and the TMM transport domain are separate.

## Materials and Methods

### Bacterial strain and growth conditions

All mycobacterial strains were grown in 7H9 (BD, Sparks, MD) medium supplemented with 0.2% glycerol, 0.05% Tween 80, and ADC (5g/L albumin, 2 g/L dextrose, 0.85 g/L NaCl, 0.003 g/L catalase). Hdb media was made as described (51) or with glycerol omitted for the soft starvation experiments. For plating the *Msmeg* strain, LB Lennox plates were used. Three different *E. coli* strains were used for cloning include DH5a, TOP10, and XL1-Blue. Antibiotic concentrations for mycobacterial strain were kanamycin - 25 µg/ml; hygromycin - 50 µg/ml; nourseothricin – 20 µg/ml; zeocin - 20 µg/ml. For *E. coli*, kanamycin – 50 µg/ml; nourseothricin - 40 µg/ml; zeocin - 25 µg/ml were used.

### Growth curves

Logarithmic phase cultures were used to perform growth curve experiments. First, three biological replicates of each strain were inoculated in 7H9 with proper antibiotics, and grown to logarithmic phase OD_600_ = 0.3, then they were diluted in 7H9 without antibiotic for the growth curve experiment. The initiation OD600 in 96 well plate was 0.1. A Synergy Neo2 linear multi- Mode Reader was used to shake the plates continuously for 18 hours at 37°C, and read OD every 15 min. The data was analyzed with exponential growth equation model using GraphPad Prism (version 9.1.2).

### Strain construction

The *mmpl3* allele swap strain was made in a strain with *wag31* knocked out at its native locus and complemented at the L5 phage integrase site (28). The *msmeg_0251-mmpl3* operon was complemented at the Giles phage integrase site, and then a knockout of both genes at the native locus was made using double stranded recombineering (52). Protein localization constructs were cloned using Gibson cloning (53). Primers are listed in supplemental table 1. The MSMEG_0251 knockout, and the CRISPRI constructs to deplete MSMEG_0250 (*mmpl3*) and MSMEG_4217 (*wag31*) and MSMEG_6929 (*murJ*) and MSMEG_4227 (*murG*) were from the Mycobacterial Systems Resource collection (31), which was a kind gift from lab of Joseph Wade, Keith Derbyshire and Ei Phoophoo Aung at the Wadsworth Center.

### Cell staining

All cells were grown to be in logarithmic phase, then incubated with HADA at 10 µM (33) for 15 min and DMN-tre at 1 µg/mL for 30 min(42) at 37°C. Then, HADA and DMN-tre were washed out and the pellets were resuspended in either 7H9, HdB +0.05% tween 80 with no glycerol (soft starvation) or in PBS tyloxapol (hard starvation) before taking images. All cells were imaged live within 20 minutes of mounting.

### Microscopy

Microscopy was done on live cells mounted on HdB agarose pads for growth conditions, or on PBS agarose pads for starvation experiments. Cells were observed on a Nikon Ti-2 widefield fluorescence microscope with a Plan Apo 100X, 1.45 NA objective. Images were collected with a Photometrics Prime 95B camera. GFPmut3, msfGFP and DMN-tre were imaged using filter cube with 470/40 nm bandpass excitation filter, a 495 nm dichroic mirror, and a 525/50 nm emission filter. Mcherry was imaged using a filter cube with 5600/40 nm bandpass excitation filter, a 585 nm dichroic mirror, and a 630/70 nm emission filter. HADA was imaged a using filter cube with 350/50 nm bandpass excitation filter, a 400 nm dichroic mirror, and a 460/50 nm emission filter. The fluorescence data was extracted from microscope images using MicrobeJ (54) and MATLAB code in (28) was used to do further evaluation.

### Elongation assays

To measure the polar length within 1.5 hours, logarithmic phase cells were used. The cells stained with HADA at 10 µM (33) for 15 min, then washed with 7H9, and the pellet resuspended in 5ml of 7H9 and outgrown for 1.5 hours at 37°C. Then the cells were stained with NADA at 10 µM for 5 minutes, washed, mounted and imaged.

### Starvation assays

In starvation experiments, cultures were grown in 7H9 to logarithmic phase, and then pelleted, washed, and resuspended in either HdB 0.05% tween 80 with no glycerol (soft starvation) or in PBS with 0.05% tyloxapol (hard starvation).

### Drug treatment of cells

All drug treatments were performed on logarithmic phase cultures in 7H9 at the following concentrations: SQ109 - 5 µg/ml, CCCP - 12.5 µg/ml, 25 µg/ml, 50 µg/ml, 75 µg/ml, 100 µg/ml, and 150 µg/ml, moenomycin -100 µg/ml, and NITD349 - 5 µM and 10 µM. The cells were incubated for 1hour at 37°C before taking images.

### Antibiotic tolerance assay

The biological replicates of MmpL3 WT and MmpL3 ΔCT strains were grown in 7H9 with proper antibiotics to logarithmic phase. Then, 7H9 was removed and the cell pellets were washed with HdB 0.05% tween 80 with no glycerol (soft starvation media). The washed pellets were resuspended in 3ml of HdB 0.05% tween 80 with no glycerol to have an OD of 0.05, then they were treated with moenomycin at 100 µg/ml. Colony-forming unit (CFU) counts were done on LB Lennox plates with zeocin 20 µg/ml.

### Minimum inhibitory concentration (MIC) assay

MmpL3 WT and MmpL3 ΔCT strains were grown in 7H9 to be in logarithmic phase, then serial dilutions (10^-1^ to 10^-6^) of three biological replicates of each strain were plated on LB Lennox plates with two-fold dilutions of antibiotics spread on the plates. Samples were also spotted on plates with no antibiotic. The MIC was determined by finding the lowest concentration of drug in which there was at least a 10-fold difference in colony number compared to the no drug control.

### RNA extraction and Q-RT-PCR

The cells were grown up to logarithmic phase (OD_600_=0.6-0.7), then the RNA was extracted from three biological replicates of each strain. The OD was normalized before starting to pellet the cultures down. 20-30 ml of each strain culture were centrifuged at 4°C for 10 min at 4000 rpm.

Then, the supernatant was removed, and the pellet was resuspended in 1000 µl of fresh TRIzol (Zymo research). The cells were lysed by bead beating for 45 and 30 seconds with 5 minutes gap between. Zymogen Direct-zol RNA Miniprep Plus (catalog no. 2070) kit and the manufacturer’s protocol was used to purify RNA. RNA samples were normalized before DNase 1 treatment (Thermofisher). To measure transcript levels of *mmpl3*, Reverse Transcription PCR (RT-PCR) was conducted with the Kapa Biosystems Sybr Fast one-step qRT-PCR kit on the normalized purified RNA of three biological replicates. The QPCR primers were designed by Primer3 (Table. 3). SigA transcript levels were used as an internal control. RT-PCR was done as described (55) in a Bio-Rad CFX Connect real-time system. To analyze the data, we used the ΔΔCT method which is described in (56).

### Thin layer chromatography (TLC) of lipid extracts

The lipids were extracted as previously described in (57), with some modifications. Cells were incubated with sodium [1-14C]-acetate (final concentration 0.2µCi.ml; Perkin Elmer) for 5 hours at 37°C. 1 ml of culture at OD600=1 were pelleted and resuspended in 800 µl of single-phase chloroform-methanol-water (2:1:0.2) solution and sonicated in bath sonication with brief vortexing (3x). Then, we added more methanol and water to get the final two-phase chloroform-methanol-water (1:1:0.8) solution. To separate the phases, the samples were centrifuged at 10,000*g for 5 min, then the aqueous phase was discarded. The organic phase (bottom phase) was collected from each sample and air-dried overnight in a fume hood.

TLC was used to analyze the extracted lipids. The chamber was equilibrated with chloroform, methanol, and water (30:8:1) for 1 hour. All dried radiolabeled lipid samples were suspended in 100 µl chloroform: methanol (4:1). Equal volumes (5 µl) of samples were mixed with 5 ml of scintillation fluid, and the [14C] was measured by a Perkin Elmer BetaScout Liquid Scintillation Counter. As the samples had different amounts of radioactivity, we normalized them to the lowest sample. An equal amount of each sample was spotted on a TLC Silica gel 60 F254 plate (Sigma Aldrich). TLC plate was developed in the chamber, then air dried for 2 hours and later visualized by Storm 860 scanner (Amersham Biosciences).The ImageQuant software (Molecular Dynamics) was used to view the images.

### Chemical synthesis

SQ109 was synthesized as previously reported (58) with minimal modification of the prior report. Briefly, *N*-isoprenyl ethylene diamine (9.8 mg, 0.05 mmol, 1.0 equiv) and 2-adamantanone (9.0 mg, 0.06 mmol, 1.2 equiv) were stirred in methanol (1.5 mL) for 2 hours at 23 °C. Then, sodium borohydride (3.8 mg, 0.1 mmol, 2.0 equiv) was added slowly over 15 minutes. The headspace of the reaction was purged with argon gas and the reaction was stirred at 23 °C for 18 hours. The reaction was diluted to 15 mL total methanol and water (20 mL) was added to the reaction to quench the remaining reducing agent. The solution was extracted with ethyl acetate (3 x 50 mL) and the organic phases were combined and evaporated *in vacuo*. The crude material was purified via flash chromatography (SiO_2_) using an isocratic eluent of 88:10:2 CHCl_3_:CH_3_OH:NH_4_OH. The purified material was evaporated *in vacuo* and converted to its HCl salt with an etherial solution of HCl gas bubbled into Et_2_O to yield a white solid, 10.8 mg (53%) yield. NMR data (CDCl_3_, 500 MHz) matched those of the previous methods (58).

## Acknowledgements.

We thank Allison Fay and the lab of Michael Glickman for the mcherry- MmpL3 construct and strains. We thank Celena Gwin and the Rego lab for the msfGFP-MmpL3 localization construct. We thank Ei Phoo Phoo Aung and the labs of Joseph Wade and Keith Derbyshire for the generous gift of the MSRdb collection. This work was funded by NIH grant R01AI148917 to CCB and a STARs award from the University of Texas System to JAB.

