## Supplemental files for "MmpL3, Wag31 and PlrA are involved in coordinating polar growth with peptidoglycan metabolism and nutrient availability"

A. Genome sequencing results of suppressors of the *wag31* point mutants

| | Strain | Mutation | Gene | Type of mutation | Site of mutation |
|---|---|---|---|---|---|
| *wag31*-D7A | sup #8 | (C)5→4 | MSMEG_0251 | Deletion (Frameshift mutation) | after TSS2 |
| | sup #21 | (C)5→6 | MSMEG_0251 | insertion (Frameshift mutation) | after TSS2 |
| | sup #22 | (GACATCGCTGTC)1→2 | MSMEG_0251 | Duplication | between TSS1 and TSS2 |
| | sup #52 | Δ1 bp | MSMEG_0251 | Deletion (Frameshift mutation) | between TSS1 and TSS2 |
| | sup #63 | W182* (TGG→TAG) | MSMEG_0251 | Missense mutation | between TSS1 and TSS2 |
| *wag31*-K20A | sup #24 | R224P (CGG→CCG) | MSMEG_0251 | Missense mutation | after TSS2 |
| | sup #33 | R224P (CGG→CCG) | MSMEG_0251 | Missense mutation | after TSS2 |
| | sup #36 | S41W (TCG→TGG) | MSMEG_0251 | Missense mutation | before TSS1 |
| *wag31*-L34A | sup #2 | (GACATCGCTGTC)1→2 | MSMEG_0251 | Duplication | between TSS1 and TSS2 |
| | sup #6 | D127G (GAC→GGC) | MSMEG_0251 | Missense mutation | between TSS1 and TSS2 |
| | sup #7 | (C)5→6 | MSMEG_0251 | insertion (Frameshift mutation) | after TSS2 |
| | sup #10 | D151G (GAC→GGC) | MSMEG_0251 | Missense mutation | between TSS1 and TSS2 |

B. Growth rate of *wag31* suppressor vs *wag31* mutants

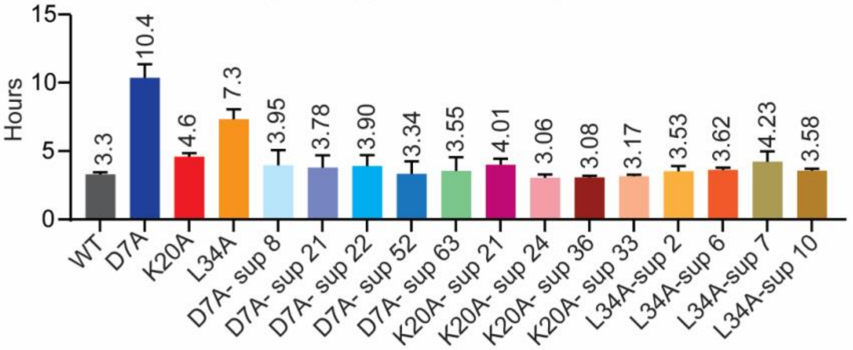

C. schematic of double-complemented strains

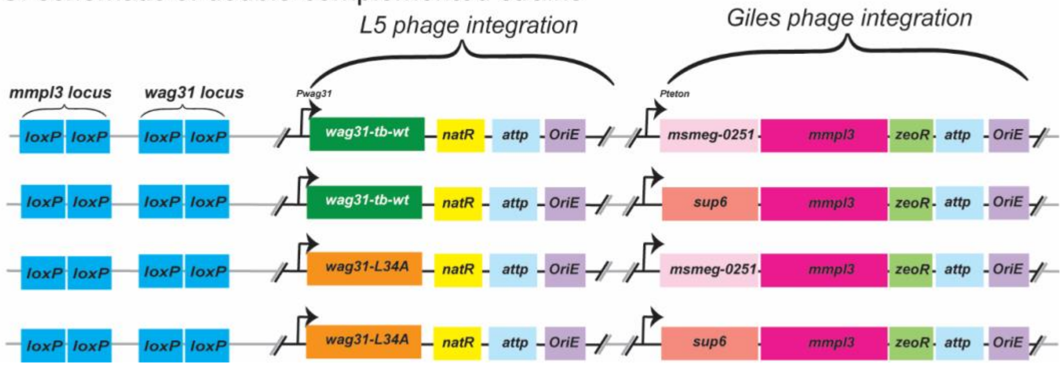

**Supplemental Figure 1.** (A) List of mutations in the *MSMEG_0251-mmpl3* operon identified in whole genome sequencing for suppressors of *wag31* point mutants. (B) Doubling time of *wag31* suppressors and parent strains. (C) The schematic of double complemented strain made by using double stranded recombineering (van Kessel and Hatfull, 2007) and used in Fig. 1B.

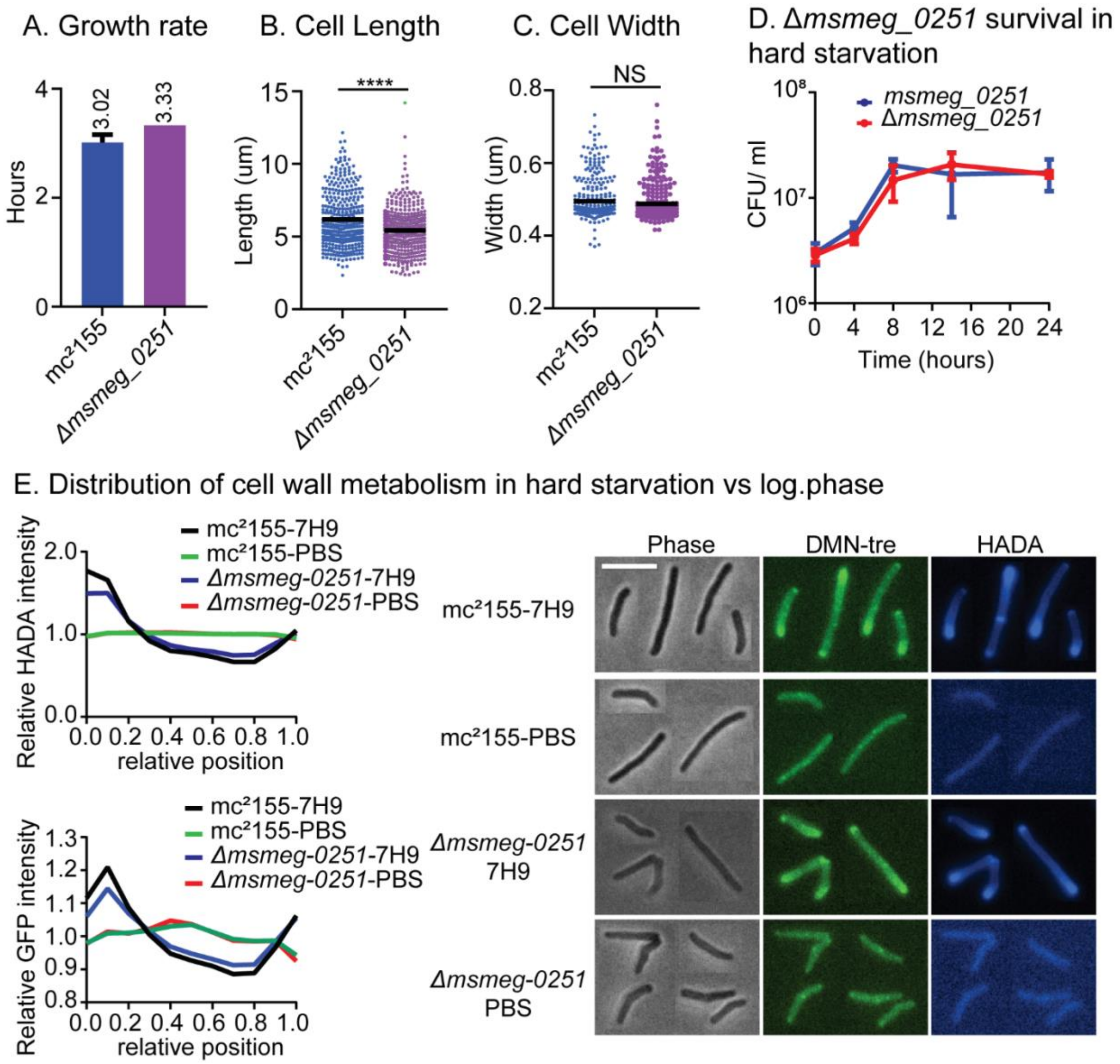

**Supplemental Figure 2. Phenotypes of ∆*msmeg_0251* strain.** (A) Doubling time of *Msmeg* cells in presence and absence of *msmeg_0251*gene. (B) and (C) The length and width in mc$^2$155 vs ∆*msmeg_0251* strain. (D) Colony Form Unit (CFU) of the expressed *msmeg_0251* (Blue line) and *msmeg_0251* deletion (Red line) strains in PBS without 0.05% Tyloxapol. (E) The cell wall metabolism in mc$^2$155 vs ∆*msmeg_0251* strain. The graphs show the distribution of peptidoglycan (top, HADA) and mycolic acid metabolism (bottom, DMN-tre) as a function of cell length, in mc$^2$155 and ∆*msmeg_0251* strain in hard starvation (PBS+ Tyloxapol) and logarithmic phase (7H9). Phase (left), DMN-tre (middle), and HADA (right) are representative images for each condition.

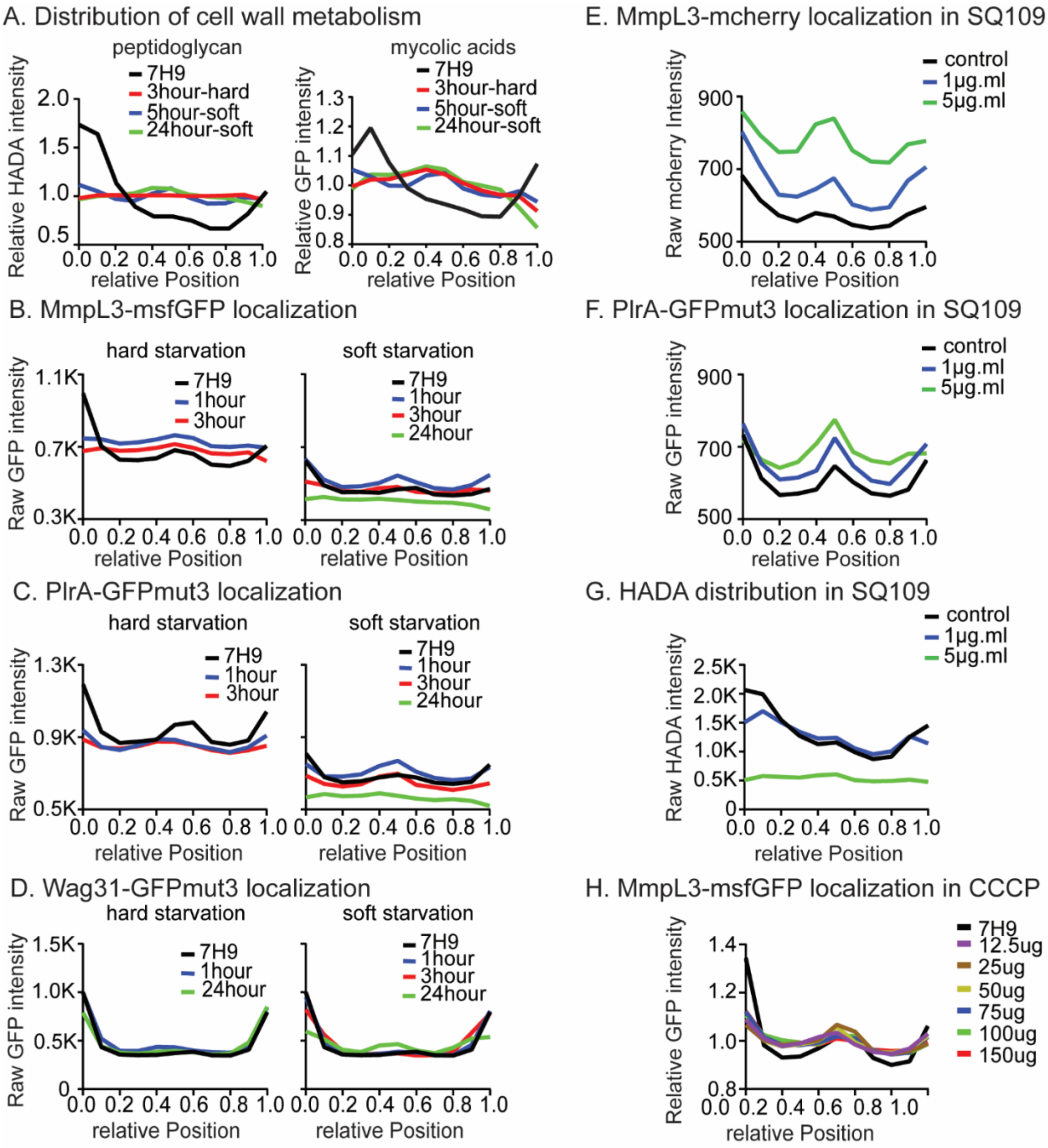

**Supplemental Figure 3. Localization of MmpL3, PlrA and Wag31 responds to nutrient availability and energy stresses.** (A) Relative, averaged HADA intensity and DMN-tre intensity, in both hard and soft starvation, from the experiment in Fig.4A. Data are oriented as in Fig.4A (B) Averaged MmpL3-msfGFP intensity in both hard and soft starvation from the experiment in Fig.4B. Data are oriented as in Fig.4B. (C) Averaged PlrA-GFPmut3 intensity in both hard and soft starvation, from the experiment in Fig.4C. Data are oriented as in Fig.4C. (D) Averaged Wag31-GFPmut3 intensity in both hard and soft starvation, from the experiment in Fig.4D. Data are oriented as in Fig.4D. (E) Averaged MmpL3-mCherry intensity from the experiment in Fig.4E. Data are oriented as in Fig.4E. (F) Averaged PlrA- GFPmut3 intensity from the experiment in Fig.4F. Data are oriented as in Fig.4F.  (G) Averaged HADA intensity of the cells with 1µg/ml and 5µg/ml of SQ109 for 1 hour. (H)  Relative, averaged MmpL3-msfGFP intensity across the length of 300+ cells treated with 12.5µg/ml, 25µg/ml, 50µg/ml, 75µg/ml, 100µg/ml, and150µg/ml for 1 hour. Data are oriented as B.
